# Auditory beat stimulation modulates memory-related single-neuron activity in the human medial temporal lobe

**DOI:** 10.1101/2020.08.27.268045

**Authors:** M Derner, L Chaieb, G Dehnen, TP Reber, V Borger, R Surges, BP Staresina, F Mormann, J Fell

**Affiliations:** Department of Epileptology, University Hospital Bonn, Venusberg-Campus 1, 53127 Bonn, Germany; Faculty of Psychology, Swiss Distance University Institute, Ueberlandstr. 12, 3900 Brig, Switzerland; Department of Neurosurgery, University Hospital Bonn, Venusberg-Campus 1, 53127 Bonn, Germany; School of Psychology & Centre for Human Brain Health, University of Birmingham, Birmingham, B15 2TT, United Kingdom

**Keywords:** Long-term memory, microwire recordings, item recognition, source recognition, binaural beats, monaural beats

## Abstract

Auditory beats are composed of two sine waves using nearby frequencies, which can either be applied as a superposed signal to both ears or to each ear separately. In the first case, the beat sensation results from hearing an amplitude-modulated signal (monaural beat). In the second case, it is generated by phase-sensitive neurons in the brain stem (binaural beat). We investigated the effects of monaural and binaural 5 Hz beat stimulation on neural activity and memory performance in neurosurgical patients performing an associative recognition task. Previously, we had reported that these beat stimulation conditions modulated memory performance in opposite directions. Here, we analyzed data from a patient subgroup, in which microwires were implanted in the amygdala, hippocampus, entorhinal cortex and parahippocampal cortex. We identified neurons responding with firing rate changes to binaural versus monaural 5 Hz beat stimulation. In these neurons, we correlated the differences in firing rates for binaural versus monaural beats to the memory-related differences for remembered versus forgotten items and associations. In the left hemisphere for these neurons, we detected statistically significant negative correlations between firing rate differences for binaural versus monaural beats and remembered versus forgotten items/associations. Importantly, such negative correlations were also observed between beat stimulation-related firing rate differences in the baseline window and memory-related firing rate differences in the poststimulus windows. In line with concepts of homeostatic plasticity, we interpret our findings as indicating that beat stimulation is linked to memory performance via shifting baseline firing levels.

## 1. Introduction

Auditory beat stimulation is a non-invasive brain stimulation technique for which effects on anxiety and cognition including memory have been reported (for overviews, see e.g. Garcia-Argibay et al. 2019, Chaieb et al. 2015). Auditory beats are amplitude-modulated tones with modulation frequencies in the range of typical electroencephalographic (EEG) rhythms. For instance, beat signals can be constructed by superposing two sine waves with nearby frequencies. Beat stimulation is either applied by presenting amplitude-modulated beat signals to one ear or both ears (monaural beats), or by presenting the original sine waves separately to each ear (binaural beats). In this latter more frequently investigated case, beat perception results from the responses of phase-sensitive brain stem neurons of the superior olivary complex (Wernick and Starr, 1968).

Based on intracranial EEG (iEEG) recordings in presurgical epilepsy patients, we demonstrated in a previous study that monaural and binaural beat stimulation caused changes in iEEG power and phase synchronization (Becher et al., 2015). These effects were most prominent at a modulation frequency of 5 Hz and were observed in temporal regions, as well as in mediotemporal structures (rhinal cortex and hippocampus), which play a crucial role in long-term memory (e.g. Eichenbaum 2000). In a subsequent study, we therefore investigated the impact of monaural and binaural 5 Hz beat stimulation on long-term memory performance in a task comprising learning and recognition of words and associated colors or scenes (Derner et al. 2018). We observed a linear effect indicating that compared to control stimulation, binaural 5 Hz beats increased and monaural 5 Hz beats decreased item memory for words, as well as source memory for associated information. These behavioral effects corresponded to reverse iEEG phase shifts within rhinal cortex for binaural versus monaural beat stimulation. However, it still remains an open question whether monaural and binaural 5 Hz beat stimulation causes changes in neural firing, which are related to the modulation of memory performance.

To answer this question, we analyzed the activity of single neurons recorded from the amygdala, hippocampus, entorhinal cortex and parahippocampal cortex in a subgroup of the previously investigated patients (Derner et al. 2018). Following spike detection and sorting, we identified neurons responding with firing rate changes to binaural versus monaural 5 Hz beat stimulation. In these neurons, we correlated the differences in firing rates for binaural versus monaural beats to the memory-related differences for remembered versus forgotten items/associations. Specifically, we assessed two alternative possibilities. Both are based on the assumption that remembering and forgetting are dependent upon the difference between memory-related and baseline firing rates: A) Beat- and memory-related firing rate changes are positively correlated: This would mean that beat stimulation-related modulations of firing rates are linked to memory performance via directly adding to memory-related firing rate changes. B) Beat- and memory-related firing rate changes are negatively correlated. This would imply that beat stimulation is linked to memory performance via shifting baseline firing levels and thereby modulating differences between memory-related and baseline firing rates. The latter possibility is in line with the Bienenstock-Cooper-Munro (BCM) learning rule (Bienenstock et al. 1982). The BCM rule proposes a sliding threshold for the induction of either long-term potentiation or long-term depression in response to alterations in neural activity. More specifically, according to this rule high/low levels of previous neuronal activity favor synaptic depression/facilitation by increasing/decreasing the crossover threshold between long-term potentiation and depression. The idea that this mechanism enables continuous adaptation of synaptic strengths to a physiological range is supported by empirical evidence (see e.g. Keck et al. 2017). Furthermore, we hypothesized that correlations between beat- and memory-related firing rate changes may in particular be observed in the left hemisphere because of its specialization for language processing (e.g. Josse and Tzourio-Mazoyer 2004).

## 2. Material and methods

### 2.1. Participants

Recordings from five presurgical epilepsy patients (3 female (age: 26/42/47 years), 2 male (age: 36/48 years)) with implanted microwires were analyzed. These patients represent a subset of a group of 13 patients (microwires had only been implanted in this subset), for whom results from macro-electrode recordings were previously reported (Derner et al., 2018). All patients gave informed written consent, and the study was conducted according to the Declaration of Helsinki, and approved by the ethics committee of the Medical Faculty of the University of Bonn.

### 2.2. Experimental paradigm

Subjects were asked to perform an associative memory task (see Figure 1) as described in an earlier study (e.g. Staresina et al., 2012). During the encoding phase of the task, 50 German nouns (per run; on each run different nouns) were presented together with a color patch (red/blue) or an image of a scene (office/nature) for 3.5 seconds each. During a jittered inter-trial interval of 700-1300 ms (mean=1000 ms) a fixation cross was presented. Subjects were asked to indicate with a button press whether the association between the color/scene and noun was plausible or not. The retrieval phase started after a 1-minute break. The 50 nouns previously presented during the encoding phase were shown together with 25 novel, previously unstudied nouns. The response options were 1) “new”, 2) the two possible color/scene sources (indicating an “old” response with source memory) and 3) a question mark (indicating an “old” response without source memory). Each response trial was displayed for a maximum of 5 seconds. In each experimental run only one source category (either color or scene) was used.

**Figure 1.**
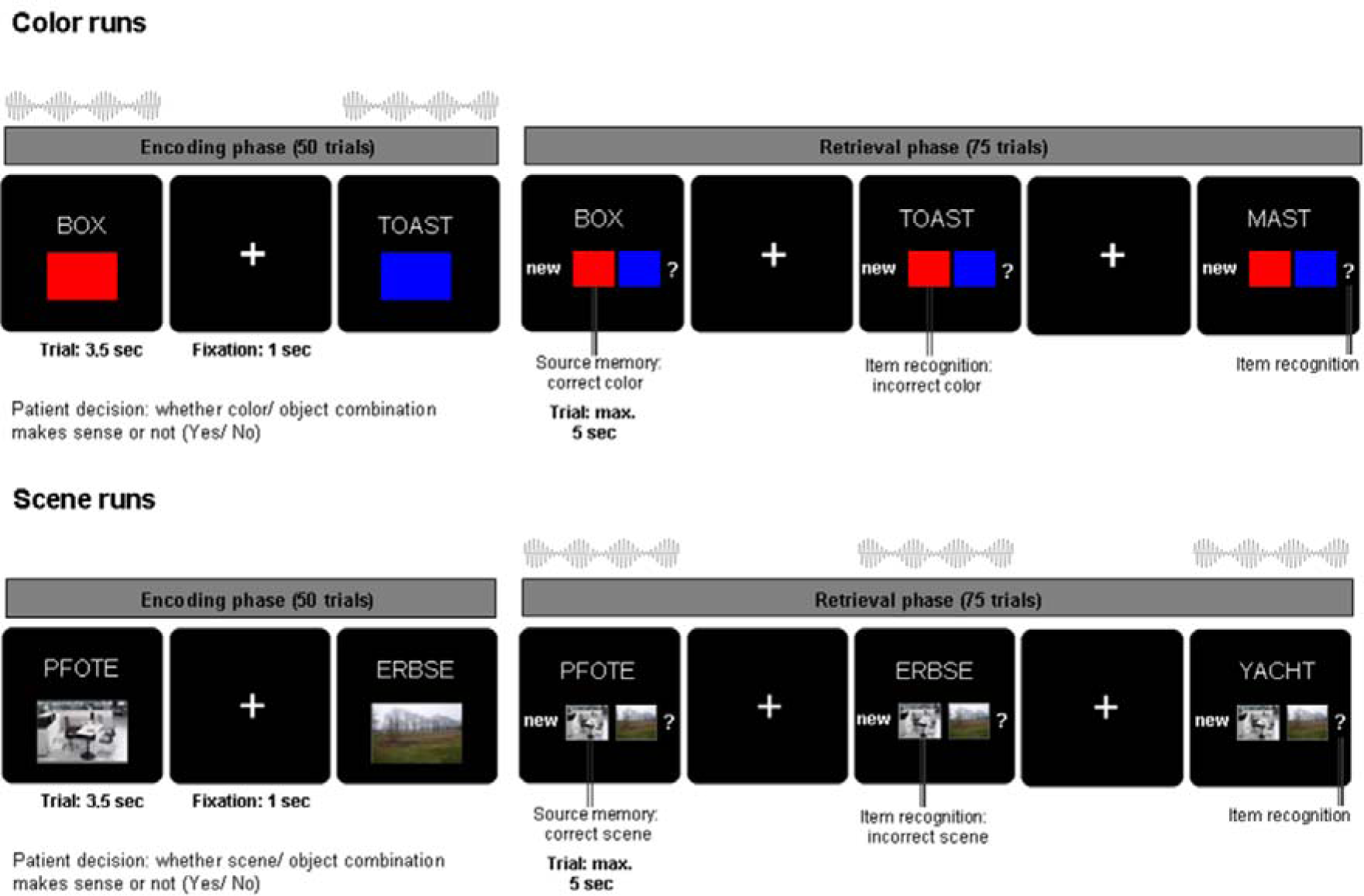
Experimental paradigm: Schematic depiction of the associative memory paradigm and the beat stimulation interventions (see Methods).

Across six experimental runs, auditory beat stimuli or a control tone were presented to the subjects either during the encoding phase (for color source runs only) or the retrieval phase (for scene source runs only) (see also Derner et al., 2018). The stimulation conditions were: binaural beats (5 Hz), monaural beats (5 Hz), control tone (220 Hz – no beat). Beat stimulation was presented at stimulus onset for the duration of each trial (encoding: 3.5s, retrieval: 5s). Auditory beats were composed of two sine waves with frequencies of 217.5 Hz and 222.5 Hz. Monaural beats resulting from the physical superposition of the two sine waves were presented to both ears simultaneously. In the case of binaural beats, one sine wave (e.g. 217.5 Hz) was presented to one ear, while the other sine wave (e.g. 222.5 Hz) was presented to the opposite ear. Each of the five subjects completed six runs each comprising 50 encoding trials and 75 retrieval trials (i.e. 300 encoding and 450 retrieval trials per subject). The order of beat stimulation conditions across the six runs had been counterbalanced within the original group of 13 patients.

### 2.3. Data recording and preprocessing

Action potential recordings were obtained from a bundle of nine microwires (eight high-impedance recording electrodes, one low-impedance reference, AdTech, Racine, WI) protruding from the end of each depth electrode targeting the hippocampus, entorhinal cortex, amygdala and parahippocampal cortex. The differential signal from the microwires was amplified using a Neuralynx ATLAS system (Bozeman, MT), filtered between 0.1 and 9,000 Hz, and sampled at 32 kHz. These recordings were stored digitally for further analysis. The number of recording microwires per patient ranged from 80 to 96. Signals were band-pass filtered between 300 and 3,000 Hz. Spike detection and sorting was performed using the Combinato software package (Niediek et al., 2016). After automated sorting using standard parameters, clusters in every channel were manually adjusted based on cluster shape, cross correlograms, and other features provided by the Combinato package. Sorted units were classified as single units, multi-units, or artifacts based on spike shape and variance, ratio between spike peak value and noise level, the inter-spike interval distribution of each cluster, and presence of a refractory period for the single units (Mormann et al. 2011).

Anatomical localization of microwires was determined based on the post-implantation CT scan co-registered to the pre-implantation MRI scan, both normalized to MNI space with SPM12. In detail, first the end of the corresponding depth electrode was visually identified. Then, microwire loci were inferred from a 3-mm-radius sphere placed 4 mm medial to the electrode tip (Staresina et al. 2019). Anatomical regions within the MTL were delineated visually according to segmentation protocols described by Insausti et al. (1998) and Pruessner et al. (2000, 2002). Only data from microwires unambiguously localized within amygdala, hippocampus, entorhinal cortex or parahippocampal cortex were kept for further analysis.

### 2.4. Evaluation of firing rates

Firing rates for each trial were calculated for four non-overlapping 500 ms intervals from 0 ms to 2000 ms relative to stimulus onset. Only units with a minimal average firing rate of 2 Hz across trials in at least one interval in one of the three stimulation conditions (in encoding or retrieval) were kept for further analysis, leaving 180 units (left side: 112, right side: 68; 95 single units, 85 multi units) from four brain regions (49 in the amygdala (27%; left: 31, right: 18), 32 in the hippocampus (18%; left: 13, right: 19), 63 in entorhinal cortex (35%; left: 49, right: 14), 36 in parahippocampal cortex (20%; left: 19, right: 17)).

### 2.5. Relation between beat stimulation and memory performance: behavioral data

In a previous study (Derner et al. 2018) we reported enhanced item and source memory for binaural compared to monaural beats during both encoding and retrieval in a group of 13 patients. For the present study, a subgroup of five patients in whom microelectrode data had been recorded, were analyzed. To evaluate the influence of beat stimulation on item and source memory in this subgroup, we conducted two-way repeated-measures ANOVAs (factors: beat condition (binaural, monaural); task phase (encoding, retrieval)) with percentage of hits and percentage of correct source decisions as dependent variables.

### 2.6. Relation between beat stimulation and memory performance: neural data

To investigate the interrelation between the behavioral effect of beat stimulation on memory and neural activity patterns (Derner et al. 2018) we first identified units which showed different firing rates for binaural vs. monaural beat stimulation separately during encoding and retrieval, as well as separately for the left and right hemisphere (Wilcoxon tests for each of the four 500 ms time windows; Bonferroni-corrected threshold of p<=0.0125). Binomial tests with probability 0.05 (corresponding to the alpha level of 5%) were conducted to test if the number of units showing a significant contrast were significantly higher than expected by chance. The significance of overlap between units related to different contrasts was calculated accordingly as portion_class1*portion_class2 (instead of 5% for single classes).

For these units we then extracted firing rates for each time window during encoding and retrieval depending on whether items (words) or sources (colors/scenes) were remembered or forgotten. This means item-related responses were classified as (later) remembered versus (later) forgotten if old words were correctly classified as old versus wrongly classified as new. Source-related responses were classified as (later) remembered versus (later) forgotten if colors/scene were correctly identified versus wrongly assigned or unknown. We then calculated the normalized firing rate (fr) differences related to beat stimulation: (fr(bin) – fr(mon))/(fr_[0;2s]_(bin) + fr_[0;2s]_ (mon)), as well as related to memory: (fr(rem) – fr(forg))/(fr_[0;2s]_(rem) + fr_[0;2s]_(forg)). Firing rate differences were normalized to exclude trivial influences of firing rate magnitude. Memory-related differences were calculated across all experimental runs in order to achieve robust estimates. As an additional analysis memory-related differences were extracted only from the runs during which beat stimulation had been applied (encoding/color and retrieval/scene runs). Finally, for each time window during encoding and retrieval, Pearson’s correlations between normalized beat stimulation-related and memory-related firing rate differences were calculated. Correlation values with a p-values below 0.0125 (Bonferroni-correction for four time windows) were considered significant.

## 3. Results

### 3.1. Behavioral data

Across the group of five subjects, the percentage of hits (correct old decisions) for color/scene runs was: 86/83% ± 5/4% (mean ± s.e.m). The percentage of false alarms (old decisions in case of new words) was: 19/19% ± 5/4%. The probability measure hits minus false alarms revealed that recognition memory was significantly above chance: 67/64% ± 4/6% (T-tests, p=0.00006/0.0004, t_4_ =17.62/10.99). The percentages of correct, incorrect, and unsure source decisions were: 63/68% ± 3/6%, 24/23% ± 7/8% and 12/9% ± 6/5%, respectively. Probability for source recognition (correct minus incorrect source decisions) was also significantly above chance: 39/45% ± 8/13% (p=0.009/0.028, t_4_ =4.68/3.37).

### 3.2. Relation between beat stimulation and memory performance: behavioral data

A two-way repeated-measures ANOVA (factors: beat condition (binaural, monaural); task phase (encoding, retrieval)) revealed a main effect of beat condition for source memory (F_1,4_ = 8.095; p < 0.05; enhanced source memory for binaural vs. monaural) and no effects or interaction for item memory; Figure 2).

**Figure 2.**
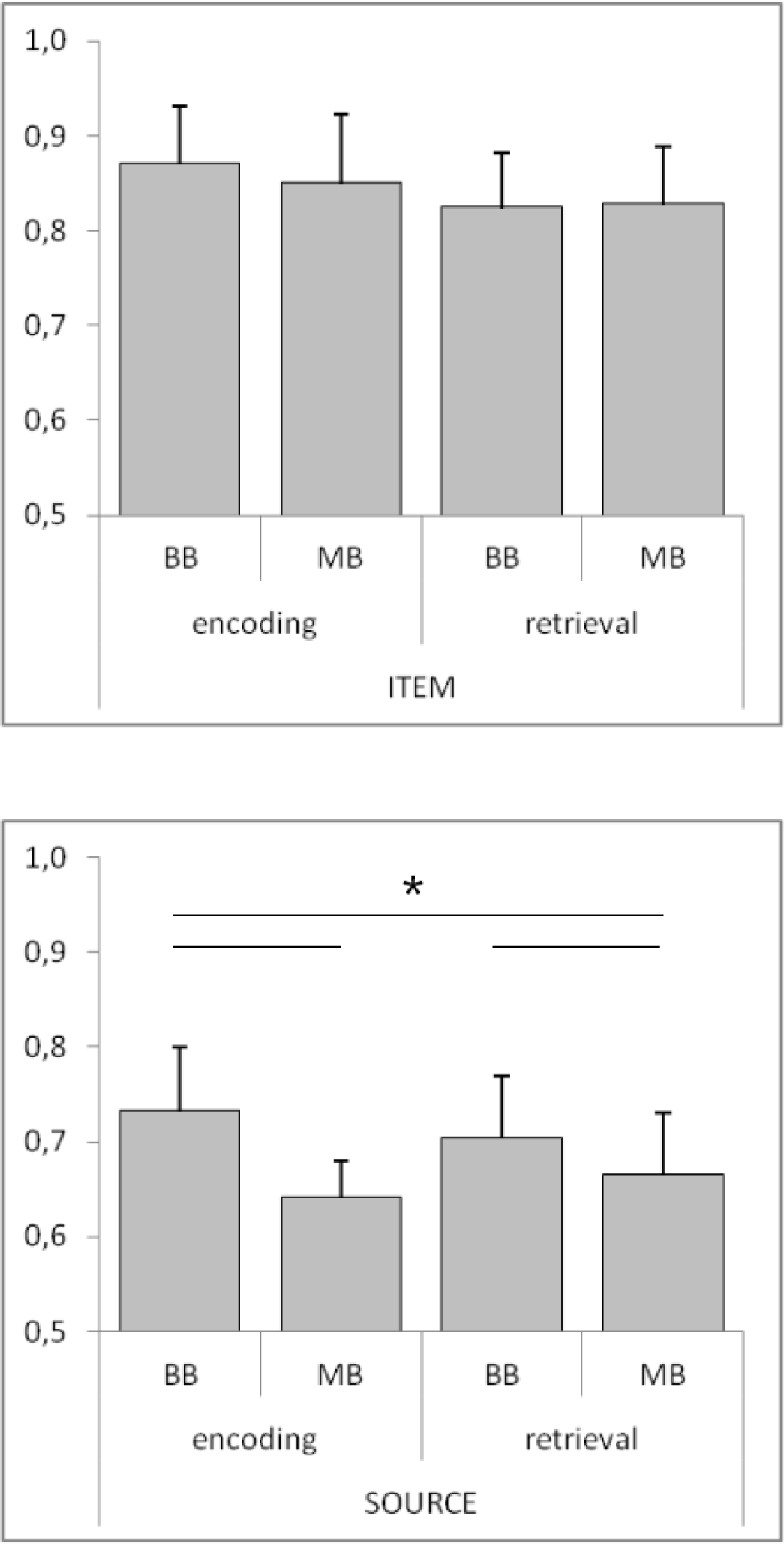
Modulation of memory performance by beat stimulation: Percentage of hits (top, item memory) and percentage of correct source responses (bottom, source memory) for binaural (BB) vs. monaural (MB) beat stimulation during encoding and retrieval. Horizontal lines and asterisks indicate a significant ANOVA main effect of beat condition (but not of task phase) for source memory (p < 0.05).

### 3.3. Relation between beat stimulation and memory performance: neural data

First, we identified neurons showing different firing rates for binaural vs. monaural beat stimulation (see Table 1). This analysis revealed 41 units for the encoding phase (23% of all units; 20 increase; 21 decrease), of these 30 on the left side (8 amygdala (26% of all amygdala units on the left side), 5 hippocampus (38%), 14 entorhinal cortex (29%) and 3 parahippocampal cortex (16%)) and 11 on the right side (4 amygdala (22%), 2 hippocampus (11%), 2 entorhinal cortex (14%) and 3 parahippocampal cortex (18%)). 75 units were found for the retrieval phase (42% of all units; 32 increase; 43 decrease), of these 52 on the left side (18 amygdala (58% of all amygdala units on the left side), 6 hippocampus (46%), 17 entorhinal cortex (35%) and 11 parahippocampal cortex (58%)) and 23 on the right side (5 amygdala (28%), 2 hippocampus (11%), 6 entorhinal cortex (43%) and 10 parahippocampal cortex (59%)). Total numbers of beat-stimulation responsive neurons were significantly higher than expected by chance both, on the left (encoding: p_binom_=3e-14; retrieval: p_binom_=3e-37; the scientific notation “e-n” stands for “10^−n^”) and right side (encoding: p_binom_=5e-4; retrieval: p_binom_=1*e-13; see table 1 for region-specific significance values). As a trend, a larger proportion of neurons differentially responded to binaural vs. monaural stimulation on the left compared to the right side (χ^2^-Tests; encoding: χ^2^_1,N=180_ = 2.71; p = 0.100; retrieval: χ^2^_1,N=180_ = 2.77; p = 0.096). Interestingly, there was no significant overlap between beat-stimulation responsive neurons during encoding and retrieval neither on the left side (17 units; p_binom_ = 0.226), nor on the right side (5 units; p_binom_ = 0.315).

**Table 1.** Overview of region-specific results of binomial tests: Results of binomial tests assessing whether numbers of neurons showing different firing rates for binaural vs. monaural beat stimulation were significantly higher than expected by chance. P-values below .05 are considered statistically significant. The scientific notation “e-n” stands for “10^−n^”. The proportions (%) of neurons with respect to the total number of neurons in each region are listed in parentheses.

|  |  | Amygdala | Hippocampus | Entorhinal cortex | Parahippocampal cortex |
| --- | --- | --- | --- | --- | --- |
| **ENCODING** | **LEFT** | 1e-04 (26%) | 3e-04 (38%) | 8e-8 (29%) | n.s. (16%) |
|  | **RIGHT** | 0.011 (22%) | n.s. (11%) | n.s. (14%) | n.s. (18%) |
| **RETRIEVAL** | **LEFT** | 4e-16 (58%) | 2e-5 (46%) | 1e-10 (35%) | 3e-10 (58%) |
|  | **RIGHT** | 0.002 (28%) | n.s. (11%) | 3e-5 (43%) | 1e-9 (59%) |

We then calculated Pearson’s correlations between normalized beat stimulation-related and memory-related firing rate differences for these neurons. There were no statistically significant correlations for the right hemisphere. However, we found several significant negative correlations for the left hemisphere (see Table 2 and Figure 3). Numerically, all but one of the 16 correlation values for the left hemisphere were negative. During encoding item memory-related firing rates were correlated with beat-related firing rates in the 1500-2000 ms window (r = -0.47, p = 0.009). During retrieval item memory-related firing rates were correlated with beat-related firing rates in the 500-1000 ms and 1000-1500 ms window (r = -0.49, p = 0.0002 and r = -0.64, p = 4*10^−7^). The latter correlation would even survive Bonferroni-correction across all tests (n = 32) conducted for both hemispheres. Finally, source memory-related firing rates during retrieval were correlated with beat-related firing rates in the 1000-1500 ms window (r = -0.38, p = 0.006).

**Table 2.** Overview of Pearson’s correlations between beat stimulation-related (BB-MB) and memory-related (REM-FORG) firing rate differences in the poststimulus windows: Correlation values for the left hemisphere are listed. Only correlations with p-values below 0.0125 (Bonferroni-correction for 4 time windows) are considered statistically significant.

|  |  | TIME WINDOW |  |  |  |
| --- | --- | --- | --- | --- | --- |
|  |  | 0-500 | 500-1000 | 1000-1500 | 1500-2000 |
| **ENCODING** | **item** | n.s. | n.s. | n.s. | -0.47 (.009) |
|  | **source** | n.s. | n.s. | n.s. | n.s. |
| **RETRIEVAL** | **item** | n.s. | -0.49 (.0002) | -0.64 (4E-07) | n.s. |
|  | **source** | n.s. | n.s. | -0.38 (.006) | n.s. |

**Figure 3.**
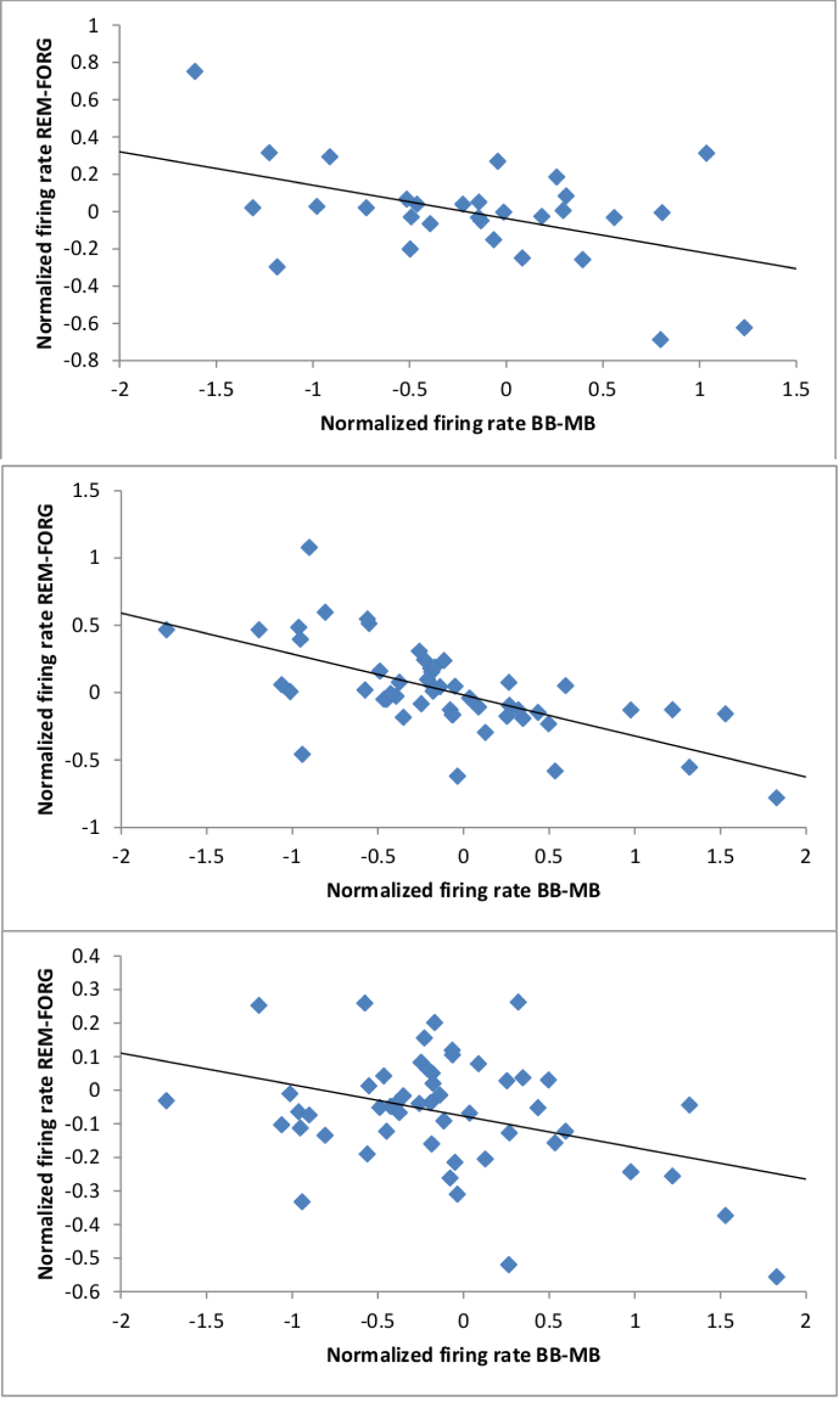
Correlation between beat stimulation-related and memory-related firing rate differences on the left hemisphere: Top: Encoding phase; item memory; 1500-2000 ms window; r = -0.47 (p = 0.009). Middle: Retrieval phase; item memory; 1000-1500 ms window; r = -0.64 (p = 4*10^−7^). Bottom: Retrieval phase; source memory; 1000-1500 ms window; r = -0.38 (p = 0.006).

Since the observed negative correlations suggest that beat stimulation has an impact on memory via shifting baseline firing levels, we further aimed to explore this possibility. Thus, we calculated Pearson’s correlations between normalized beat stimulation-related firing rate differences in the baseline window [-500ms;0ms] and memory-related firing rate differences in the poststimulus windows for the same neurons (see Table 3). Corresponding to the four cases of statistically significant negative correlations described above three time windows again showed significant negative correlations. Moreover, this analysis revealed two additional cases of negative correlations. Beat stimulation-related firing rate differences in the baseline window were negatively correlated with memory-related firing rate differences in the 1000-1500 ms window for source encoding (r = -0.47, p = 0.008), as well as in the 0-500 ms window for item retrieval (r = -0.36, p = 0.010). These findings corroborate the idea that beat stimulation-related shifts of baseline firing levels mediate the effects on memory performance.

**Table 3.** Overview of Pearson’s correlations between beat stimulation-related (BB-MB) firing rate differences in the baseline window [-500;0ms] and memory-related (REM-FORG) firing rate differences in the poststimulus windows: Correlation values for the left hemisphere are listed. Only correlations with p-values below .0125 (Bonferroni-correction for 4 time windows) are considered statistically significant.

| normiert mit mean 0-2000 |  | TIME WINDOW |  |  |  |
| --- | --- | --- | --- | --- | --- |
|  |  | 0-500 | 500-1000 | 1000-1500 | 1500-2000 |
| **ENCODING** | **item** | n.s. | n.s. | n.s. | -0.47(0.009) |
|  | **source** | n.s. | n.s. | -0.47(0.008) | n.s. |
| **RETRIEVAL** | **item** | -0.36 (0.010) | -0.48 (0.0003) | -0.67 (4E-08) | n.s. |
|  | **source** | n.s. | n.s. | n.s. | n.s. |

Finally, we performed an additional analysis based on extracting memory-related differences only for the actual runs with beat stimulation (encoding/color and retrieval/scene runs). In line with the previous analyses, we observed a significant negative correlation between beat-related and item memory-related firing rates during retrieval in the 1000-1500 ms window (r = -0.38, p = 0.006). A negative correlation was also evident between beat-related baseline firing rates and item memory-related firing rates in this time window (r = -0.43, p = 0.002). Moreover, we found a positive correlation between beat-related and source memory-related firing rates during retrieval in the 0-500 ms window (r = 0.36, p = 0.009). However, there was no significant correlation (r = 0.31, p = 0.025) between beat-related baseline firing rates and source memory-related firing rates in this time window.

## 4. Discussion

An impact of beat stimulation on memory performance has been described by several groups (for overviews, see e.g. Garcia-Argibay et al. 2019; Chaieb et al. 2015). Recently, we reported that binaural and monaural 5 Hz beat stimulation modulated memory performance in opposite directions based on data from a larger patient group (Derner et al. 2018). Binaural beats were related to enhanced and monaural beats to impaired item and source memory. In the subgroup analyzed here, only a significant behavioral effect on source memory was evident. In accordance with previous intracranial EEG findings (Becher et al. 2015, Derner et al. 2018), we observed a significant effect of binaural versus monaural beat stimulation on firing rates in a large fraction of neurons within the medial temporal lobe. As a trend, a larger proportion of neurons differentially responded to beat stimulation on the left versus right side. Similarly, a recent study reported that theta EEG responses to 6 Hz binaural beats were dominant in the left hemisphere (Jirakittayakorn and Wongsawat 2017). Interestingly, numbers of beat-stimulation responsive neurons in parahippocampal cortex were significantly above chance during retrieval, but not during encoding (table 1). This outcome may be related to the fact that parahippocampal neurons were shown to play an important role in scene processing (Mormann et al. 2017) and that stimulation during retrieval occurred during scene runs, while stimulation during encoding occurred during color runs.

For the beat-stimulation responsive neurons, we detected statistically significant negative correlations between firing rate differences for binaural versus monaural beats and remembered versus forgotten items/associations in the left hemisphere. Importantly, such negative correlations were also observed between beat stimulation-related firing rate differences in the baseline window and memory-related firing rate differences in the poststimulus windows. Therefore, we interpret our findings as indicating that beat stimulation is linked to memory performance via shifting baseline firing levels and not via directly adding to memory-related firing rate changes. Expressly speaking, we suggest that by shifting baseline levels differences between memory-related and baseline firing rates are modulated consistent with increased memory performance for binaural versus monaural beats.

Our results are in accordance with the BCM rule of homeostatic plasticity (Bienenstock et al. 1982), proposing that high levels of previous neuronal activity favour synaptic depression and low levels favour facilitation. Experimental support for such an interrelation has, for instance, been reported in studies investigating the impact of light deprivation on visual processing in rats (Kirkwood et al. 1996), as well as transcranial direct current stimulation on learning of motor behavior (Bortoletto et al. 2015) and visuomotor coordination in humans (Antal et al. 2008). Moreover, a compensatory increase/decrease in synaptic strength after chemically or optogenetically induced inhibition/excitation of neural activity has also been shown in rodent hippocampal neurons (Lee et al. 2013; Mendez et al. 2018). Finally, the left-hemispheric localization of our findings is in line with its well-known specialization for language processing (Josse and Tzourio-Mazoyer 2004). Taken together, our results support the hypothesis that auditory beat stimulation has an impact on memory performance via modulation of single neuron activity within the medial temporal lobe.

## Conflicts of interest

The authors declare that no financial or non-financial competing interests exist.

## Acknowledgements

This work was supported by the German Research Foundation (DFG SFB 1089, MO 930/8-1).

## Notes

### Competing Interest Statement

The authors have declared no competing interest.

